# “Identification and enrichment of SECReTE *cis*-acting RNA elements in the *Coronaviridae* and other (+) single-strand RNA viruses”

**DOI:** 10.1101/2020.04.20.050088

**Authors:** Gal Haimovich, Tsviya Olender, Camila Baez, Jeffrey E. Gerst

## Abstract

*cis*-acting RNA motifs play a major role in regulating many aspects of RNA biology including posttranscriptional processing, nuclear export, RNA localization, translation and degradation. Here we analyzed the genomes of SARS-CoV-2 and other single-strand RNA (ssRNA) viruses for the presence of a unique *cis* RNA element called SECReTE. This motif consists of 10 or more consecutive triplet nucleotide repeats where a pyrimidine nucleotide (C or U) in present every third base, and which we identified in mRNAs encoding secreted proteins in bacteria, yeast, and humans. This motif facilitates mRNA localization to the endoplasmic reticulum (ER), along with the enhanced translation and secretion of translated protein. We now examined for SECReTE presence in Group IV and V RNA viruses, the former including the *Coronaviridae*, like SARS-CoV-2 and other positive (+)ssRNA viruses, and the latter consisting of negative (-) ssRNA viruses. Interestingly, the SARS-CoV-2 genome contains 40 SECReTE motifs at an abundance of ~1.3 SECReTEs/kilobase (kb). Moreover, all ssRNA viruses we examined contain multiple copies of this motif and appears in (+)ssRNA viruses as non-random in occurrence and independent of genome length. Importantly, (+)ssRNA viruses (*e.g.* Coronaviruses and Hepaciviruses), which utilize ER membranes to create double membrane vesicles to serve as viral replication centers (VRCs), contain more SECReTE motifs per kb as compared to (−)ssRNA viruses (*e.g*. Rabies, Mumps, and Influenza), that replicate in the nucleus or the cytoplasm, or other (+)ssRNA viruses (*e.g*. Enteroviruses and Flaviviruses) which employ different organellar membranes. As predicted by our earlier work, SECReTE sequences are mostly found in membranal or ER-associated/secreted proteins. Thus, we propose that SECReTE motifs could be important for the efficient translation and secretion of secreted viral proteins, as well as for VRC formation. Future studies of SECReTE function and identification of SECReTE-binding proteins could provide new drug targets to treat COVID-19 and other (+)ssRNA related diseases.

## Introduction

Human infection with *Coronaviridae* (CoV) viruses can result in severe acute respiratory distress leading to lethality. The recent outbreak of the SARS-CoV-2 virus emphasizes the potent ability of human coronaviruses (hCoVs) to infect and rapidly spread throughout the human population, given the absence of immune prophylaxis (*e.g.* vaccination) or curative treatment. SARS-CoV-2 is a positive (+) single-strand RNA [(+)ssRNA] virus comprising a genome of ~30kb encoding at least 29 viral proteins (VPs) involved in viral infection, replication, and release^1,2^. The genome is organized into a 5’untranslated region (UTR)-leader sequence, followed by a large open reading frame (ORF1ab) that encodes 16 non-structural VPs (nsp1-16), then by ORFs encoding the viral accessory proteins and structural proteins (*e.g.* spike (S), envelope (E), membrane (M), nucleocapsid (N)), and terminating with a 3’UTR-polyA tail. The non-structural VPs are involved in the cleavage of polypeptide1ab (NSP5), suppression of host antiviral response (nsp1), creation of the viral replication center from the endoplasmic reticulum (ER) (nsp2,3,4,6), and viral RNA replication (nsp7,8,9,10,12,13,14,15,16). Four structural VPs (S,E,M,N) form the coat of the virus and along with other small ORFs facilitate virion assembly, release, and infection. As with other (+)ssRNA viruses, upon infection the SARS-CoV-2 RNA acts as an mRNA for the direct translation of viral ORF1ab.

As with other members of the hCoVs, infection of the lung epithelia with SARS-CoV-2 likely induces a reticulo-vesicular network of ER-derived double membrane vesicles (DMVs) that form a discrete viral replication organelle (or center; VRC)^3–5^. VPs are translated on the VRC surface, with many (*i.e.* soluble and membrane-anchored proteins) translocated into the membrane of the newly forming structure. VRC formation, therefore, represents an essential step for both vRNA replication and virion assembly, and hence, progressive infection upon virion release. Yet, little is known of the organellar dynamics and interactions with either the viral RNA or VPs to create the replication membrane, although morphological alteration of the ER (and, perhaps, other secretory pathway organelles, *e.g.* lipid droplets, Golgi, endosomes) is consistent with the secretory nature of viral replication, which first involves nsp translation and translocation.

The ER is the primary site for the translation and translocation of soluble secreted and membrane (secretome) proteins. Thus, vRNA interactions with ER-associated RNA-binding proteins (RBPs) likely constitute a critical rate-limiting step in VP production. In our work on RNA trafficking and association with intracellular organelles as a means to regulate protein translation and localization, we recently identified a *cis* RNA element present in nearly all secretome proteins from bacteria to humans^6^. This motif, entitled “<u>s</u>ecretion-<u>e</u>nhancing *<u>c</u>is* <u>r</u>egulatory <u>t</u>argeting <u>e</u>lement” (SECReTE), is based upon extended (>10 consecutive) triplet nucleotide repeats whereby a pyrimidine nucleotide is present every third base, whether in coding regions (as *NYN* or *NNY*; where *N* is any nucleotide and *Y* = U or C) or in the UTR regions. Mutational analyses performed using several yeast genes encoding secreted proteins (*e.g. SUC2*, *CCW12*, *HSP150*) revealed that the addition or removal of SECReTE motifs in yeast mRNAs could enhance or inhibit mRNA stability and association with the ER, respectively, thereby affecting protein secretion and cell physiology. Thus, SECReTE is important for mRNA-protein interactions at the level of the ER that facilitate protein production. We now identify numerous SECReTE motifs encoded in many of SARS-CoV-2 VPs, particularly in those encoding membrane-associated proteins.

Viral SECReTE motifs (vSECReTEs) were found in the genes of all ssRNA viruses inspected and its abundance (*i.e.* SECReTE/kb) correlates overall with the percentage of C and T nucleotides (%CT) in the genomes (we use “T” instead of “U” henceforth for the convenience of sequence analysis). However, when looking at specific %CT levels we see a wide variability in SECReTE score between viral families/genera. Importantly, we found that (+)ssRNA viruses (*e.g. Coronaviridae* and *Hepaciviruses*) that utilize ER-derived DMVs for replication centers^3–5,7^ contain more SECReTE motifs per kb as compared to (−)ssRNA viruses (*e.g*. *Rhabdoviridae, Orthomyxoviridae*, and *Paramyxoviridae*), which replicate in the nucleus or the cytoplasm^8–13^, or other (+)ssRNA viruses which use different organellar membranes and do not form ER-derived DMVs (e.g. *Enteroviruses, Nodavirridae*, and *Flaviviruses*)^4,5,7,14^. Interestingly, the position of some vSECReTE motifs along the length of the Spike gene of all seven human coronaviruses is quite similar. This co-occurrence may indicate the possibility of conservation/convergence of motif position. Thus, we predict that SECReTE motifs may be important for the association of viral RNA with the ER, as well as for the efficient translation of viral membrane proteins and creation of VRCs at ER membranes. Thus, continued studies of SECReTE function and the identification of SECReTE-binding proteins could provide new drug targets to treat COVID-19, and other (+)ssRNA related diseases.

## Results

### SECReTE elements are present in the human *Coronaviridae*

Because of the current COVID-19 pandemic we questioned whether SECReTE elements are present in viruses, particularly those of the hCoVs, and whether they may fulfill a role in viral replication and virion production. We first determined if SARS-CoV-2 genomic RNA (gRNA) contains SECReTE motifs using the same script we used to identify SECReTEs in yeast and human mRNAs^6^. We found that the ~30kb SARS-CoV-2 gRNA contains forty motifs (Figure 1a). All SECReTE elements are located in protein coding sequences (CDS) and 72.5% of the SECReTE elements are encoded in either membranal or secretion-associated proteins (*e.g.* nsp3, nsp4, nsp6, ORF7a, ORF7b, S, M, E and N proteins). Notably, all motifs, but one, are in *NNY* or *NYN* frames (Figure 1A & Supplementary Table S1). This result is similar to our findings in yeast and humans, in which mRNAs encoding secretome proteins that contain either a signal peptide (SP) or TMD, as well as mRNAs encoding secreted proteins that lack these domains, contain SECReTE elements^6^.

**Figure 1.**
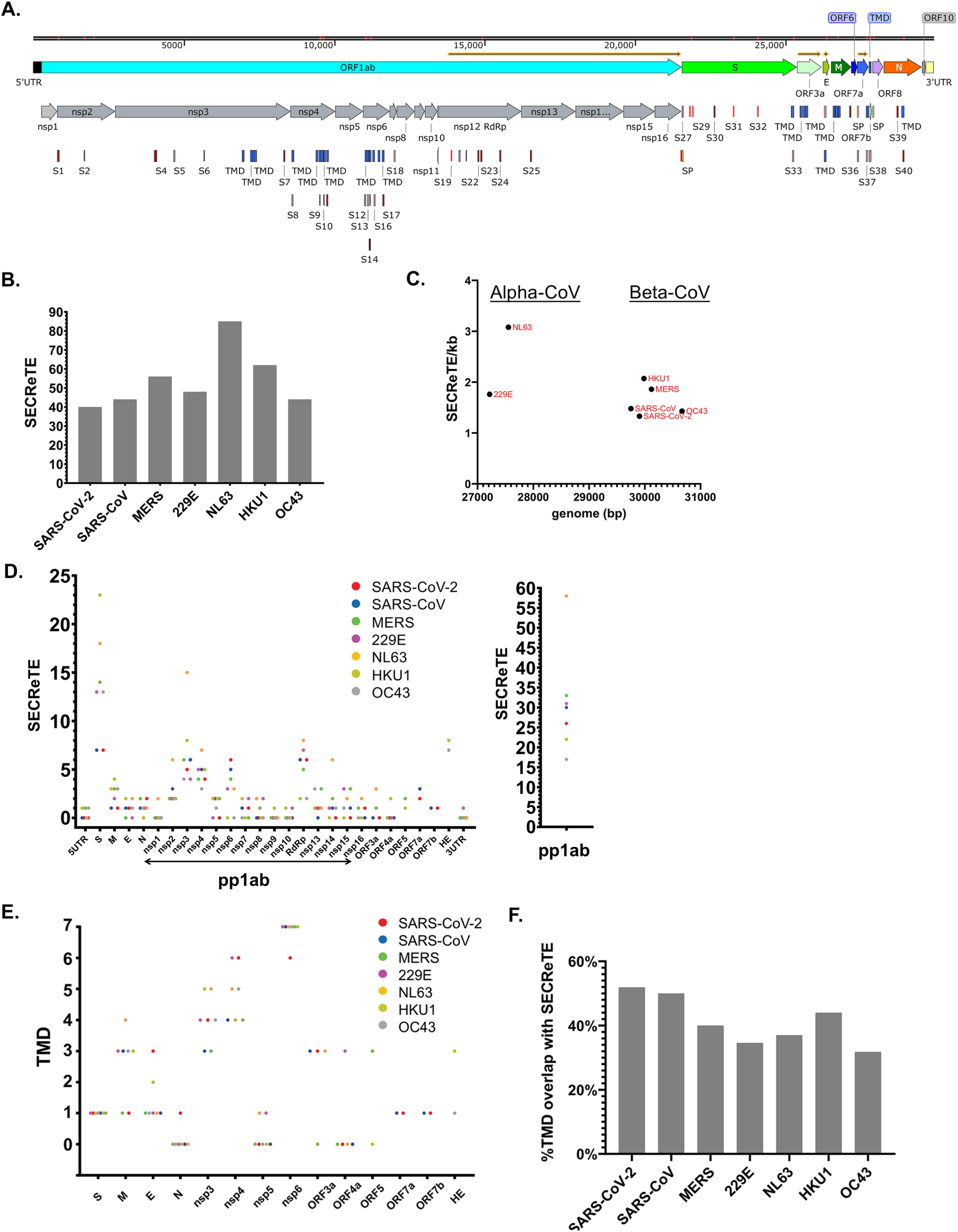
SECReTE elements in SARS-CoV-2 and other human coronaviruses. **A)** Schematic of the SARS-CoV-2 viral genome with listing of vSECReTE elements (annotated S#1-40); TMD = transmembrane domain, SP = signal peptide. **B)** A graph depicting the number of vSECReTEs in each of the human coronaviruses (hCoVs). **C)** A graph depicting the SECReTE score (SECReTE/kb) *vs* genome size of the seven hCoVs. **D)** The distribution of vSECReTE motifs along hCoVs untranslated regions (UTRs) and coding sequences. Each dot represents the number of vSECReTE of the indicated region, color-coded by hCoV species. Note that not all hCoVs have all the depicted genes. Right side: the combined number of vSECReTEs of all 16 non-structural proteins which are translated as pp1ab protein. **E)** The number of transmembrane domains (TMD) in each protein, color coded as in panel D. **F)** The percentage of TMD-encoding sequences that have a SECReTE motif, calculated for each of the 7 hCoVs. See also Supplementary Table S1.

To test whether these SECReTE sequences are common to all SARS-CoV-2 strains, we screened for the SECReTE motifs in the genomes of 493 different isolates of SARS-CoV-2 (Supplementary Table S2). We found 20 cases (~4%) in which a mutation occurred in a SECReTE motif (Table 1). In one case the mutation resulted in motif elimination, while in another case the mutation resulted in motif creation. Two cases resulted in the shortening or elongation of the motif. All other mutations, whether synonymous or non-synonymous, did not affect the existence of the motif or its length. Currently, we cannot determine if these mutations affect the pathogenicity, infectivity or other aspects of SARS-CoV-2 biology.

**Table 1.**
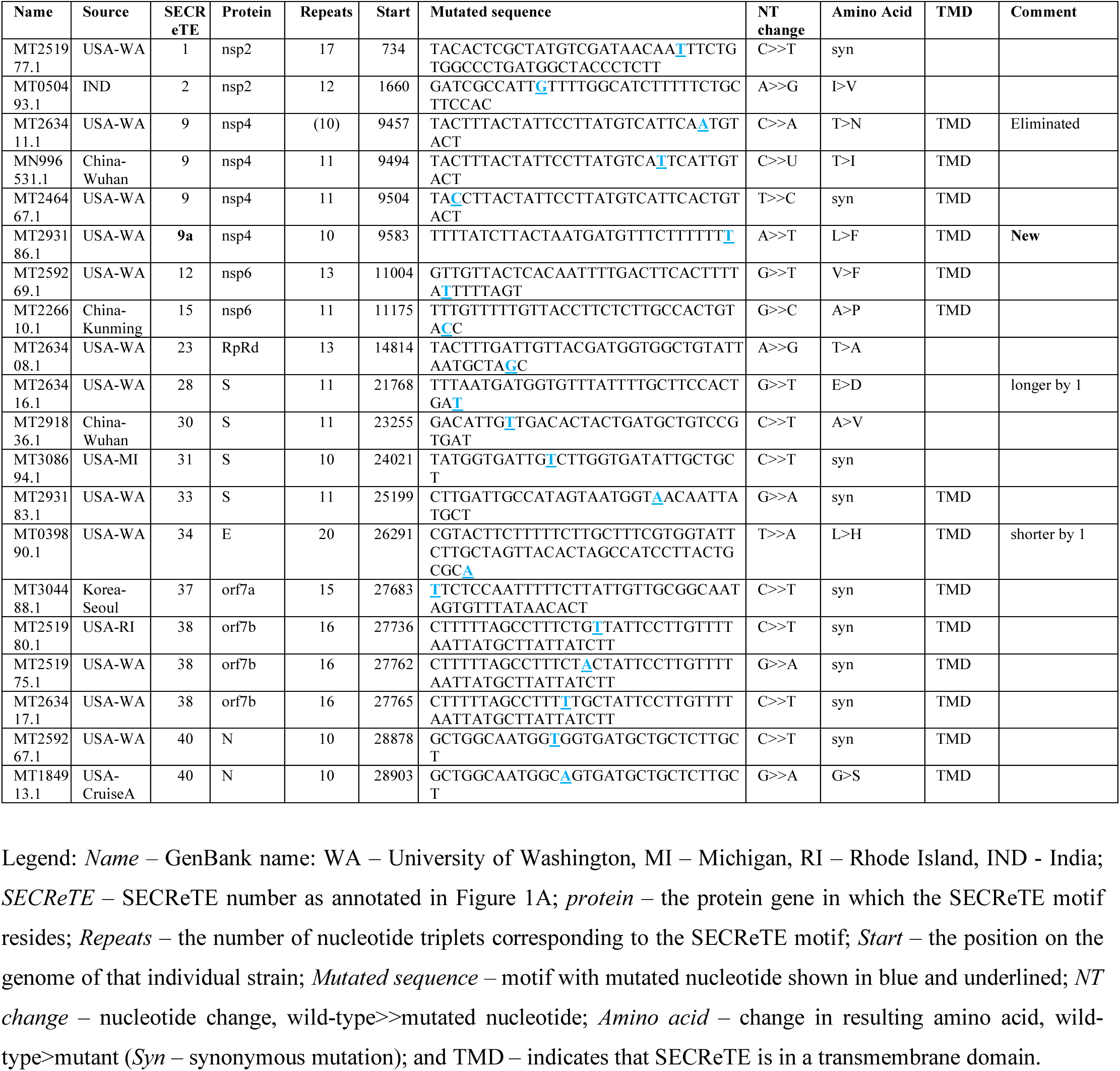
Analysis of 493 SARS-CoV-2 strains reveals mutations in SECReTE motifs

Next, we compared the SECReTE content in SARS-CoV-2 to six other human coronaviruses, *e.g.* SARS-CoV, MERS-CoV, hCoV-NL63, hCoV-229E, hCoV-OC43 and hCoV-HKU1. SARS-CoV and MERS-CoV are considered potentially pandemic coronaviruses that cause severe acute respiratory syndrome, similar to SARS-CoV-2, but are less infectious and appear to result in a higher mortality rate than SARS-CoV-2^15^. The other four viruses are endemic to the human population and typically cause mild respiratory ailments with very low mortality rates^16^. Of note, hCoV-NL63 and hCoV-229E belong to the Alpha-CoV genus with a slightly shorter genome as compared to the other five CoVs, which belong to the Beta-CoV genus. All seven viruses contain SECReTE sequences in varying amounts (*i.e.* between 40-85 SECReTE motifs), the highest being hCoV-NL63 (Figures 1B-C and Supplementary Table S1). Based upon the limited data available regarding the infectivity and lethality of the different human CoVs, we did not observe an association between the number of SECReTE sequences and viral pathogenicity when comparing the three highly pathogenic (*e.g.* SARS-CoV, SARS-CoV-2, and MERS) to the three least pathogenic (*e.g.* hCoV-229E, HKU1 and OC43), particularly if the moderately pathogenic^17^ and most SECReTE-enriched hCoV, hCoV-NL63, is not included (Supplementary Figure S1). At this juncture we cannot draw any conclusions regarding a relationship between SECReTE elements and viral infectivity, replication, or pathogenicity.

### SECReTE elements are present in human CoV genes encoding membranal or secreted proteins

As with SARS-CoV-2, SECReTE sequences are not distributed equally along the length of the other hCoV genomes, but are concentrated in specific genes. A large number of motifs (*e.g.* 4-6) are each found in the ORFs encoding the S, nsp3,4, and 6, and the RNA-dependent RNA polymerase (nsp12/RdRp) proteins (Figure 1D). Although RdRp is not a secreted protein *per se*, it does need to be translated on ER membranes and localizes to the DMV. Hence, we suspect that the high number of SECReTE motifs in the RdRp sequence may facilitate better RNA localization to, and translation at, ER membranes/DMVs. The hemaglutinin esterase (HE) gene, which is present only in the hCoV-OC43 and hCoV-HKU1 genomes, also contains a large number (*e.g.* 7-8) of SECReTE elements. Only three of the hCoVs examined have SECReTE elements in their 5’UTRs and only one has a motif in the 3’UTR. Since the non-structural proteins are translated as two long polypeptides (pp1a and pp1ab), we scored the total number of SECReTE of pp1ab for each virus. As expected, pp1ab contains the largest number of SECReTEs, with a high variability between the different CoVs. The only exception is hCoV-HKU1, which contains more SECReTE sequences in S as compared to pp1ab.

Similar to the situation in yeast and human mRNAs, SECReTE elements are mainly found in genes encoding secretome proteins [*i.e.* secreted proteins likely to possess transmembrane domains (TMDs) and/or signal peptides (SP)^6^ and Supplementary Table S3]. In SARS-CoV-2, fourteen SECReTE elements are found in TMDs out of a total of twenty-seven TMDs, and two elements were found in SPs out of total of three SPs present (Figure 1A and Supplementary Table S1). SARS-CoV-2 has the highest number of TMD sequences with SECReTE motifs out of the seven human CoVs examined (Figures 1E-F).

To test whether SECReTE sequences might be positionally conserved among the seven hCoVs, we aligned the S gene sequences and looked for co-occurrences of the SECReTE motifs. Strikingly, the SECReTE motif at the beginning of the gene (SECReTE27) is maintained in six out of seven hCoVs and SECReTE33, located in the single encoded TMD, is maintained in five out of seven hCoVs (Supplementary Figure S2A & B and Supplementary Table S1). Note that in hCoV-229E, there is no SECReTE in the TMD sequence, but there is a SECReTE motif (SECReTE44 of hCoV-229E) downstream and almost parallel in position to SECReTE33 of SARS-CoV-2. SECReTE31, located in the in the middle of the S gene, is also maintained in five of seven hCoVs (and in hCoV-229E SECReTE38 is 45nt downstream). Only one SARS-CoV-2 S gene motif (SECReTE 29) had no parallel in any of the other hCoVs. Importantly, position of the abovementioned SECReTEs is maintained even when the nucleotide and amino acid sequences differ.

### Correlation of SECReTE abundance with the %CT content of the CoV genomes

Next, we expanded our analysis to 2993 genomes of *Coronaviridae* viruses and strains that were organized by similarity to the hCoVs^18^ or by their genera (Supplementary Table S4). We plotted the SECReTE score (SECReTE number/kb) *vs.* %CT based on these two groupings and found a good correlation between the SECReTE and %CT in at least some of the groups (R^2^ = 0.6830, 0.8637, 0.5763 for Alpha-CoV, Beta-CoV, Delta-CoV, respectively) and a low correlation (R^2^ = 0.04333) with Gamma-CoV) (Figure 2A). For 229E-like, HKU1-like, NL63-like, OC43-like, MERS-like, SARS-like and SARS-2-like, respectively, the R^2^ values were 0.1794, 0.7710, 0.6801, 0.6982, 0.8759, 0.01020 and 0.003998 (Figure 2B). Notably, for some CoV families/genera we observe a wide variability in SECReTE scores that at the same %CT, which might arise via non-random occurrences (see below). Alpha-CoVs have the most SECReTEs/kb and at any %CT (Figure 2A), as compared to the other genera, with hCoV-NL63 being the most extreme (Figure 2B). Overall, nearly all CoV genomes had a SECReTE score of >1 SECReTE/kb.

**Figure 2.**
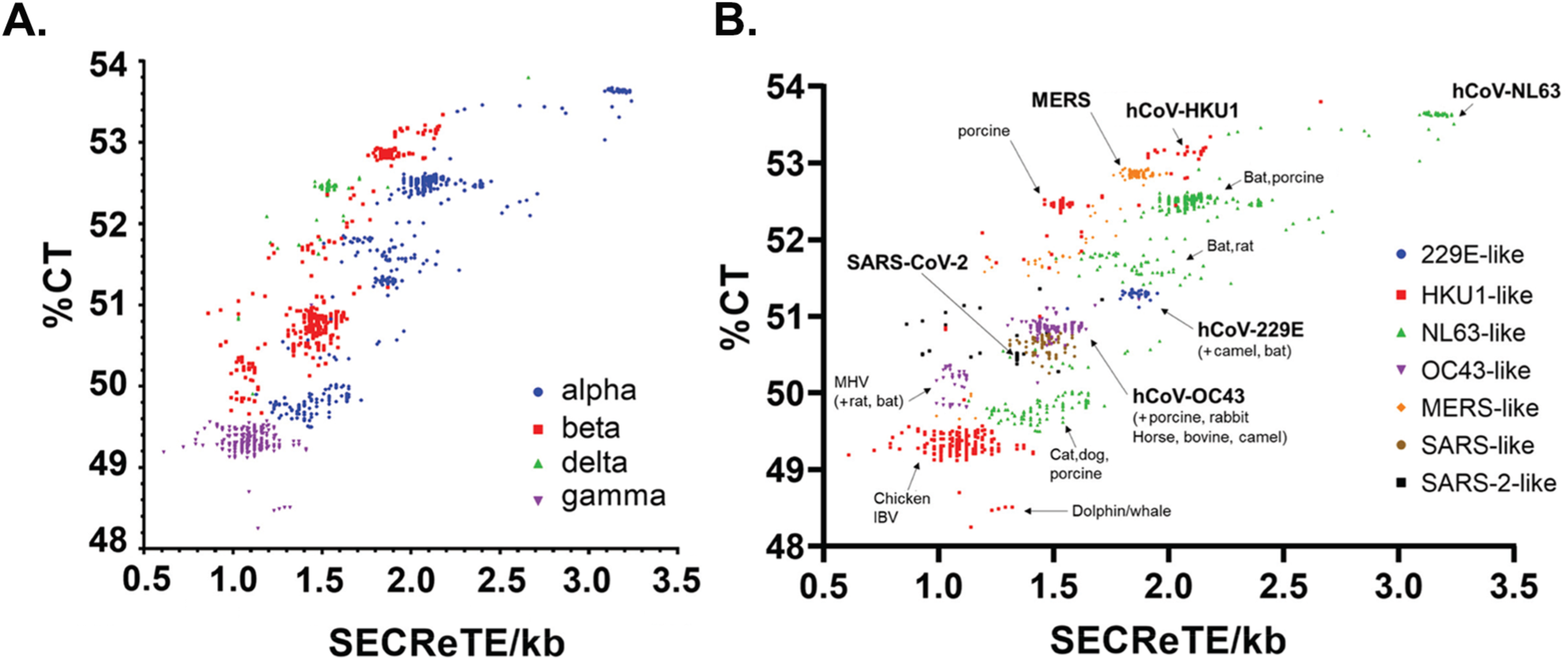
SECReTE elements in the *Coronaviridae* family. **A)** Distribution of the SECReTE score of each viral genome, color coded by Genus, *vs.* the percentage of pyrimidines (%CT) in the genome. **B)** Distribution of the SECReTE score of each viral genome, color coded by similarity to one of the 7 hCoVs^18^ *vs.* the percentage of pyrimidines (%CT) in the genome. Note in both panels that there are multiple strains of certain viruses. The position of the 7 hCoVs and CoVs infecting other animal hosts are depicted. See also Supplementary Table S4.

### (+)ssRNA viruses have a higher %CT and SECReTE content than (−)ssRNA viruses

We then identified SECReTE motifs in CDS’s from 463 (+)ssRNA, 119 (−)ssRNA, and 47 ambisense ssRNA viruses from seventeen different viral families [Figure 3A and Supplementary Tables S5 (vSECReTE scores) & S6 (vSECReTE sequences and position)]. The data shows that (−)ssRNA viruses have a lower SECReTE score as compared to (+)ssRNA viruses. Note that in this plot we scored genomic segments separately (*e.g*. for segmented genome (−)ssRNA viruses, like influenza or ambisense RNA viruses). Thus, segments that lack SECReTEs have a SECReTE score of ‘0’. To gain further insight, we averaged the SECReTE score for each family or distinct genera and plotted them *vs*. %CT (Figures 3B and Supplementary Figure S3). We found a large variability in SECReTE content amongst (+)ssRNA viral families (*e.g.* ranging from ~0.3 to ~5 SECReTE/kb). However, all (−)ssRNA and ambisense viruses examined show, on average, 0.28-0.5 SECReTE/kb. The difference between (+)ssRNA and (−)ssRNA viruses was statistically significant (p<2.2 e-16; Kolmogorov-Smirnov test for viruses with %CT between 47% to 55%). As before, we found an overall correlation to the %CT (R^2^ =0.6299). We note, however, that genome size does not correlate with the SECReTE score (R^2^ =0.01045) (Figure S4).

**Figure 3.**
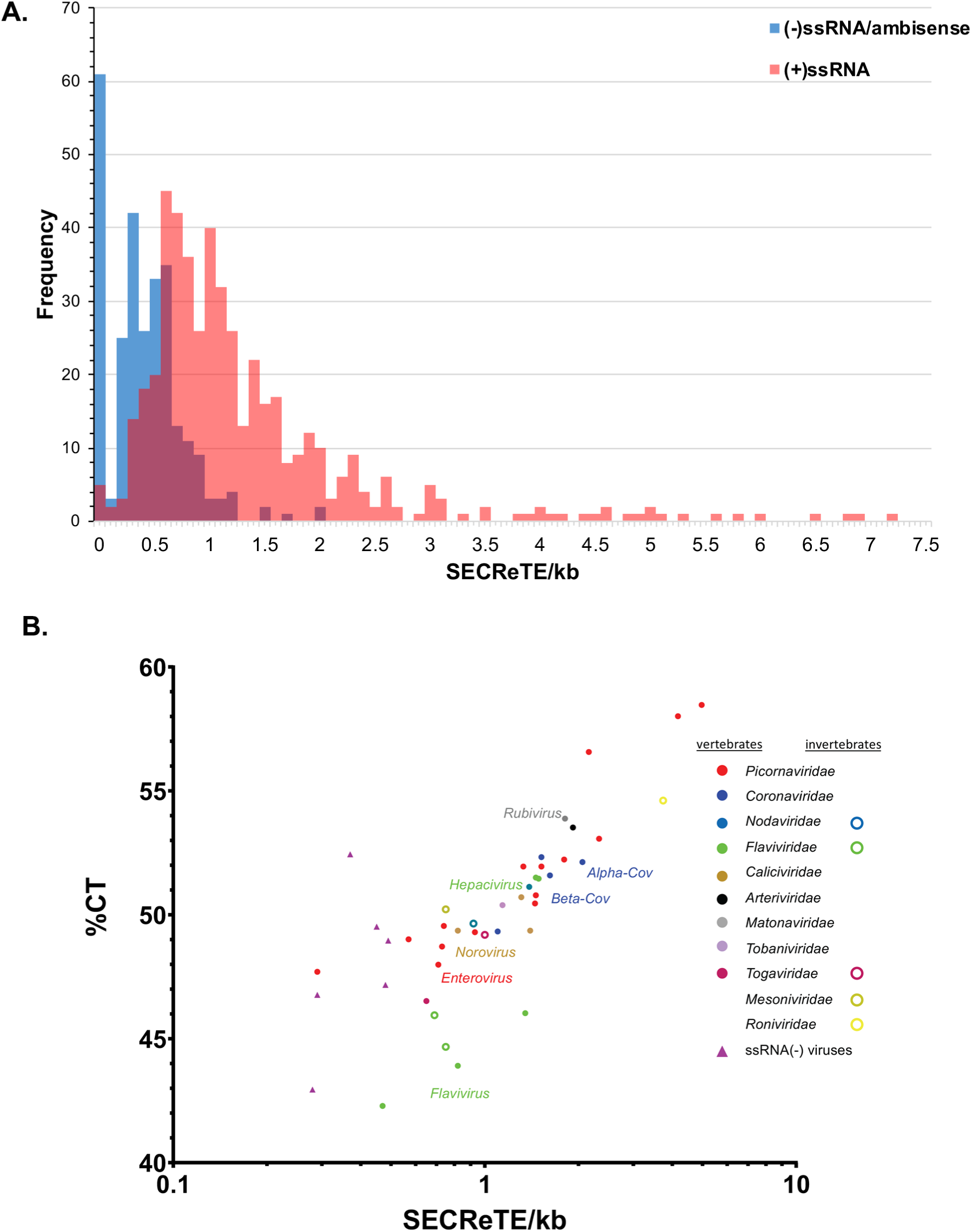
SECReTE elements in (+)ssRNA and (−)ssRNA viruses. **A)** A histogram depicting the frequency of each SECReTE score (by bins of 0.1) for all viruses tested. **B)** The distribution of the average SECReTE score of each family/genus (color-coded) *vs*. the average percentage of pyrimidines (%CT) in the genomes. Open circles – viruses that infect only invertebrates. Closed circles – viruses that infect vertebrates or use invertebrates as vectors to infect vertebrates (see also Supplementary Tables S5 & S6 and Supplementary Figure S3).

To overcome differences in the %CT of different viruses, we tested the significance of SECReTE appearance in specific viral genomes via permutation analysis by re-shuffling (500 repetitions) the viral genomes and scoring motif number after each round. By applying permutations, each virus genome is compared to sequences with exactly the same %CT. While the SECReTE score in (+)ssRNA viruses was significantly higher than expected by random chance, this was not the case for (−)ssRNA viruses (Table 2). Thus, the ubiquitous and specific presence of SECReTE elements suggests that they could play a role in RNA translation directly from (+)ssRNA genomes.

**Table 2.** The SECReTE score of (−)ssRNA viruses is lower than expected by chance

| Accession | Virus genome type | Family | Species name | Genome size | %CT | #SECRete | >wt permutation in | p value |
| --- | --- | --- | --- | --- | --- | --- | --- | --- |
| NC_001542 | negative | *Rhabdoviridae* | Rabies virus | 11932 | 48.40 | 6 | 54 | 0.10978 |
| NC_001608 | negative | *Filoviridae* | Marburg virus | 19111 | 49.08 | 13 | 15 | 0.031936 |
| NC_001796 | negative | *Paramyxoviridae* | Human parainfluenza Virus 3 | 15462 | 43.24 | 5 | 13 | 0.027944 |
| NC_002200 | negative | *Paramyxoviridae* | Mumps rubulavirus | 15384 | 48.84 | 10 | 24 | 0.0499 |
| U18101 | negative | *Rhabdoviridae* | Spring viremia of carp virus | 11019 | 44.16 | 2 | 106 | 0.213573 |
| NC_004162 | positive | *Togaviridae* | Chikungunya virus | 11826 | 45.14 | 6 | 3 | 0.007984 |
| NC_005831 | positive | *Coronaviridae* | **Human Coronavirus** NL63 | 27553 | 53.66 | 85 | 0 | 0.001996 |
| NC_010354 | positive | *Picornaviridae* | Foot-and-mouth disease virus | 8201 | 49.41 | 12 | 0 | 0.001996 |
| NC_019843 | positive | *Coronaviridae* | **MERS** coronavirus | 30119 | 52.84 | 53 | 0 | 0.001996 |
| NC_024770 | positive | *Picornaviridae* | Chicken gallivirus 1 | 8432 | 58.47 | 38 | 0 | 0.001996 |
| NC_028970 | positive | *Picornaviridae* | Theilovirus | 8101 | 54.36 | 8 | 0 | 0.001996 |
| NC_038307 | positive | *Picornaviridae* | Polio virus | 7440 | 47.35 | 5 | 0 | 0.001996 |
| NC_045512 | positive | *Coronaviridae* | **SARS-CoV-2 isolate** Wuhan-Hu-1 | 29903 | 50.45 | 40 | 0 | 0.001996 |

Legend: *Accession* – GenBank accession number; *Virus type* – ssRNA virus; *Size* – genome size in nucleotides; *%CT* – percentage of pyrimidines in viral genome; *#SECRete* – number of SECRetes in the wild-type viral genome; *>WT in permutation* – the number of permutations (out of 500) in which the sequence contained more SECRete sequences than the wild-type genome; *p value* – probability of statistical significance.

### Human mRNAs with high SECReTE scores

Because of the connection between SECReTE score and viral biology, we thought that a high SECReTE score might also be unique to certain human gene families. By screening the human secretome genes (Supplementary Table S3) we found three gene families with a particularly high SECReTE scores when compared to other genes of similar size or function (Figure 4). These gene families included the mucins, defensins, and olfactory receptors (Figure 4), known to be expressed within epithelial cells and olfactory neurons, respectively. These families scored significantly higher than most other genes (p=0.04 for defensins *vs*. a random list of similar sized genes; and p<0.0001 for comparisons of OR *vs*. GPCRs, or a random list of similar sized genes, and mucins *vs.* a list of random genes of similar size, as well as for each group *vs.* all genes; unpaired t-test).

**Figure 4.**
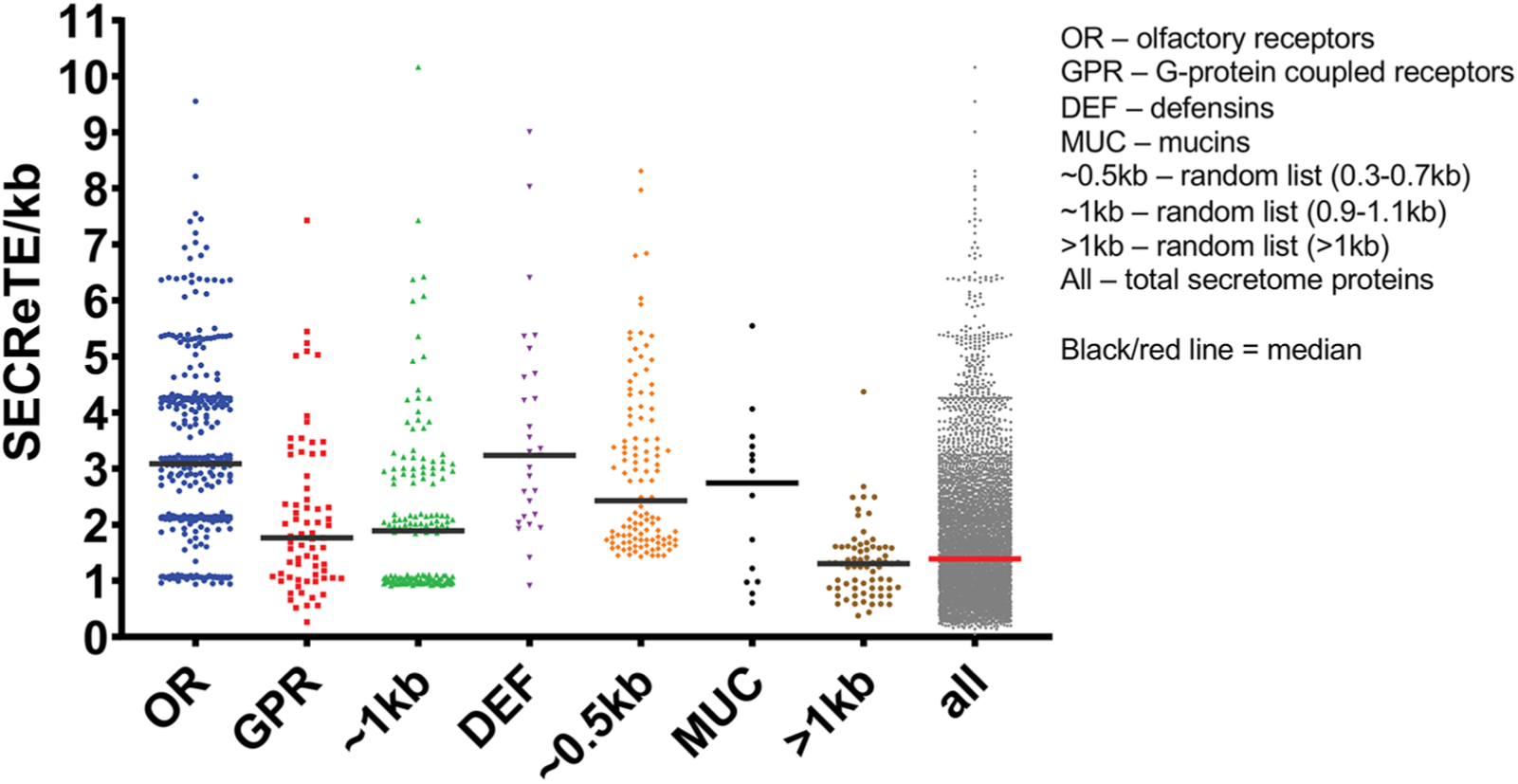
SECReTE score in human secretome genes shows high-scoring gene families. The graph shows the SECReTE score of olfactory receptor (OR), mucins and defensins mRNAs compared to the SECReTE scores of G-protein-coupled receptors, a random list of secretome genes of similar average sizes as ORs (~1kb), defensins (~0.5kb) and mucins (>1kb), and all secretome genes. Each dot is a single gene. Lines depict the medians. See also Supplementary Table S3.

## Discussion

Previously, we identified SECReTE as a pyrimidine-rich *cis*-acting RNA element prevalent in secretome-encoding genes from bacterial to human cells and demonstrated its role in facilitating protein secretion from yeast cells^6^. Here we demonstrate that SECReTE motifs are not exclusive to cells, but are also prevalent in viral genomes, including (+)ssRNA, (−)ssRNA, and ambisense RNA viruses (Supplementary Tables S1,4-6). Given the COVID-19 pandemic, we focused primarily on the (+)ssRNA viruses of the *Coronaviridae*, which includes SARS-CoV-2 (Figure 1A) and other hCoVs. Interestingly, SECReTE motifs are more enriched in the (+)ssRNA viruses, as opposed to the (−)ssRNA and ambisense RNA viruses (Figure 3A and B), and notably in the hCoVs (Figures 1B and C). SECReTE motifs are found in hCoV genes encoding either structural or non-structural viral proteins (Figure 1D) and primarily, though not exclusively, in genes for membranal and secreted viral proteins (Figures 1E and F). We also found that the level of SECReTE abundance per kb correlated well with the percentage of pyrimidines (%CT) present in the viral genomes within the (+)ssRNA and (−)ssRNA viruses in general (Figure 3). However, there was variability in this correlation within the hCoVs (Figures 2A and B) and subsequent permutation analyses revealed that SECReTE occurrence in *Coronaviridae* and (+)ssRNA viruses is probably non-random (Table 2). In addition, SECReTE abundance did not correlate with viral genome length (Figure S4).

Because of the COVID-19 pandemic, we looked for a potential correlation between the SECReTE score to viral pathogenicity (relative to SARS-CoV-2 and the other hCoVs). While the SECReTE scores of the hCoVs are largely similar, hCoV NL63 had the highest number of SECReTEs and SECRETEs per kb (Figures 1B & C). Interestingly, NL63 may be more pathogenic than the other endemic hCoVs, although this supposition is based upon a very limited number of reports totaling less than 200 cases world-wide^19–25^, and hence its inclusion in the highly pathogenic hCoV group is pending more research (Supplementary Figure S1). Overall, we could not form a definitive association between SECReTE score and hCoV pathogenicity. Interestingly, however, we could begin identify several potential phenomenon related to SECReTE presence in both viral and human genes. First, we observed that SECReTE positions within the hCoV Spike protein showed co-occurrence (Supplementary Figure S2 and Supplementary Table S1). This suggests that motif presence could be positionally conserved, although the mechanism by which this happens (*i.e.* conservation *vs.* drift *vs.* convergence) is not known. Nevertheless, it suggests a functional requirement for SECReTE in some aspect of either S gene RNA association with membranes or in Spike protein translation, or perhaps both. Further work is necessary to reveal the role (if any) of SECReTE motifs in hCoV RNA association with ER membranes or the translation of viral proteins, not only for the S gene, but for all other structural and non-structural protein-encoding genes containing this motif. Since SECReTE presence or absence significantly affected the synthesis and secretion of three yeast proteins examined (Suc2, Hsp150, and Ccw12), as well as an exogenously expressed form of secretion-competent GFP in yeast^6^, we presume that the motif may fulfill a similar role in the production of viral proteins. Second, we determined that certain families of human proteins are enriched with SECReTE motifs more than random occurrence (Figure 4). Interestingly, these families included mucins and defensins which are secreted from or associated with epithelial layers that are targeted by hCoVs. This could be coincidence or it could be that host mRNAs and hCoV vRNAs use the same SECReTE-interacting proteins expressed in these cells. While entirely speculative, it would not be improbable to think that organization of the VRC on ER membranes and usurpation of host SECReTE-binding proteins could strongly influence the translation and translocation of secreted proteins, as well as tethering the viral genome to the ER membrane and the formation of DMVs. A negative impact upon the translation of proteins involved in innate immunity (*e.g.* defensins) and barriers to infection (*e.g.* mucins) alone might be expected to render cells more sensitive to infection^27^. Interestingly, both anecdotal preliminary evidence for the onset of anosmia, which can occur upon OR or olfactory neuron loss, has been recently described for COVID-19 patients^28^. Hence the identification of host proteins that interact with hCoV viral RNA elements may prove important not only for understanding viral biology, but may also uncover druggable targets that interfere with viral propagation. Overall, our work shows that SECReTE RNA elements are present throughout all biological entities and suggests that they are likely to be important also for viral replicative cycle.

## Methods

SECReTE identification and %CT calculations were performed as previously described^6^. TMD identification was performed with TMHMM v2.0^26^. Alignment of the S gene sequence of the seven hCoVs was done using SnapGene™. Detection of overlap between SECReTE sequences and TMD or within aligned S genes was done manually. For the permutation analysis, we shuffled (500 times) the coding sequences of selected viruses genes. The statistical significance of the SECReTE count per virus was calculated as (#successes +1) / (number of permutations + 1), where #successes is the number of permuted sequences with a SECReTE count higher than the native genome. Because genes in viruses tend to overlap, redundant SECReTE sequences were merged for calculation of the SECReTE count.

## Acknowledgments

This work was supported by a grant from the Jeanne and Joseph Nissim Center for Life Sciences Research, Weizmann Institute of Science and, in part, by a grant from the Israel Science Foundation (#578/18). J.E.G. holds the Besen-Brender Chair in Microbiology and Parasitology, Weizmann Institute of Science.

---

**Supplementary Figure S1.**
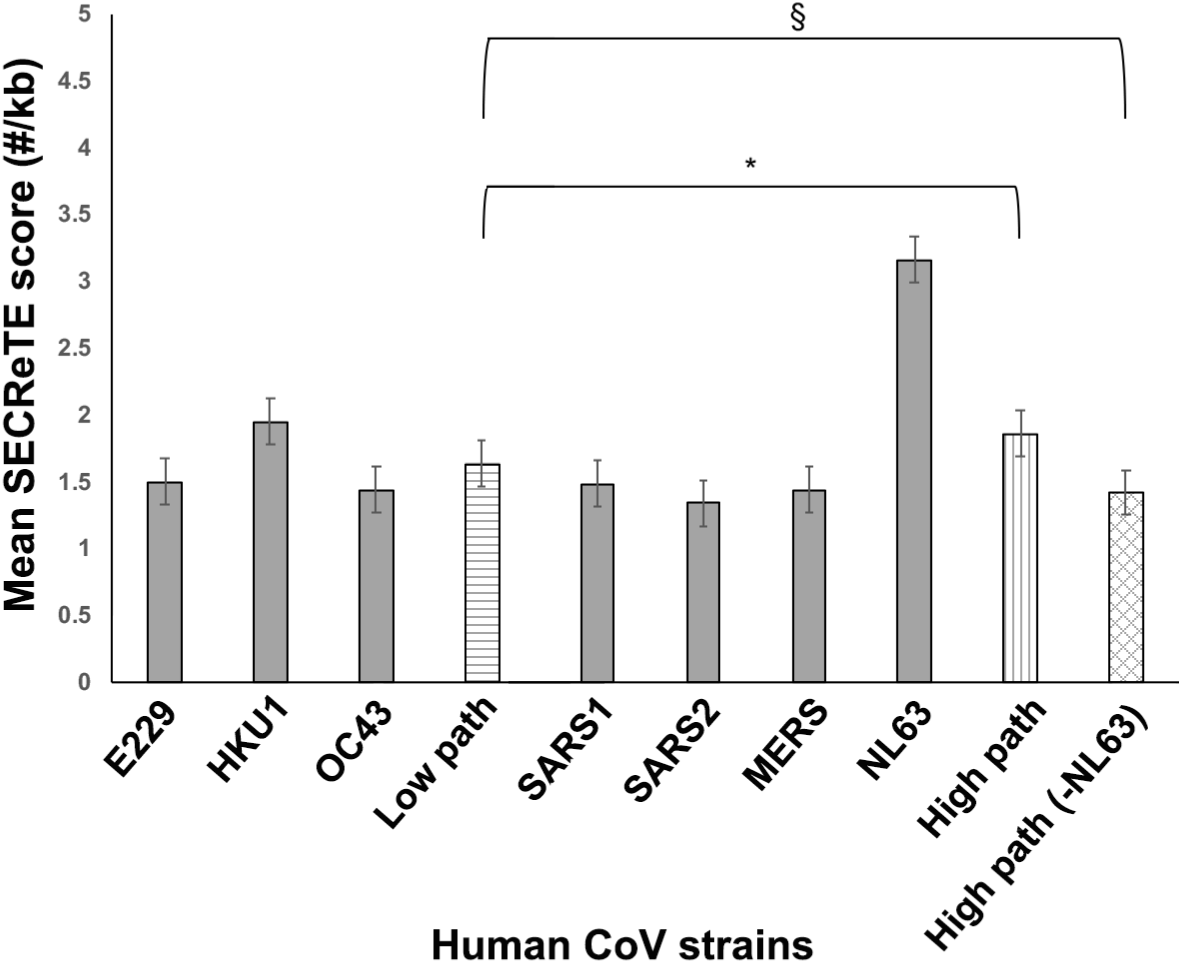
SECReTE score does not correlate with pathogenicity of hCoVs. The SECReTE scores of individual hCoV (grey bars) were averaged based on the pathogenicity (*Low Path –* low pathogenicity; *High Path –* high pathogenicity) of the viruses (high or low, longitudinal or latitudinal stripes, respectively). Checkered – high pathogenicity without NL63. * – p<0.05. § – p>0.05.

**Supplementary Figure S2.**
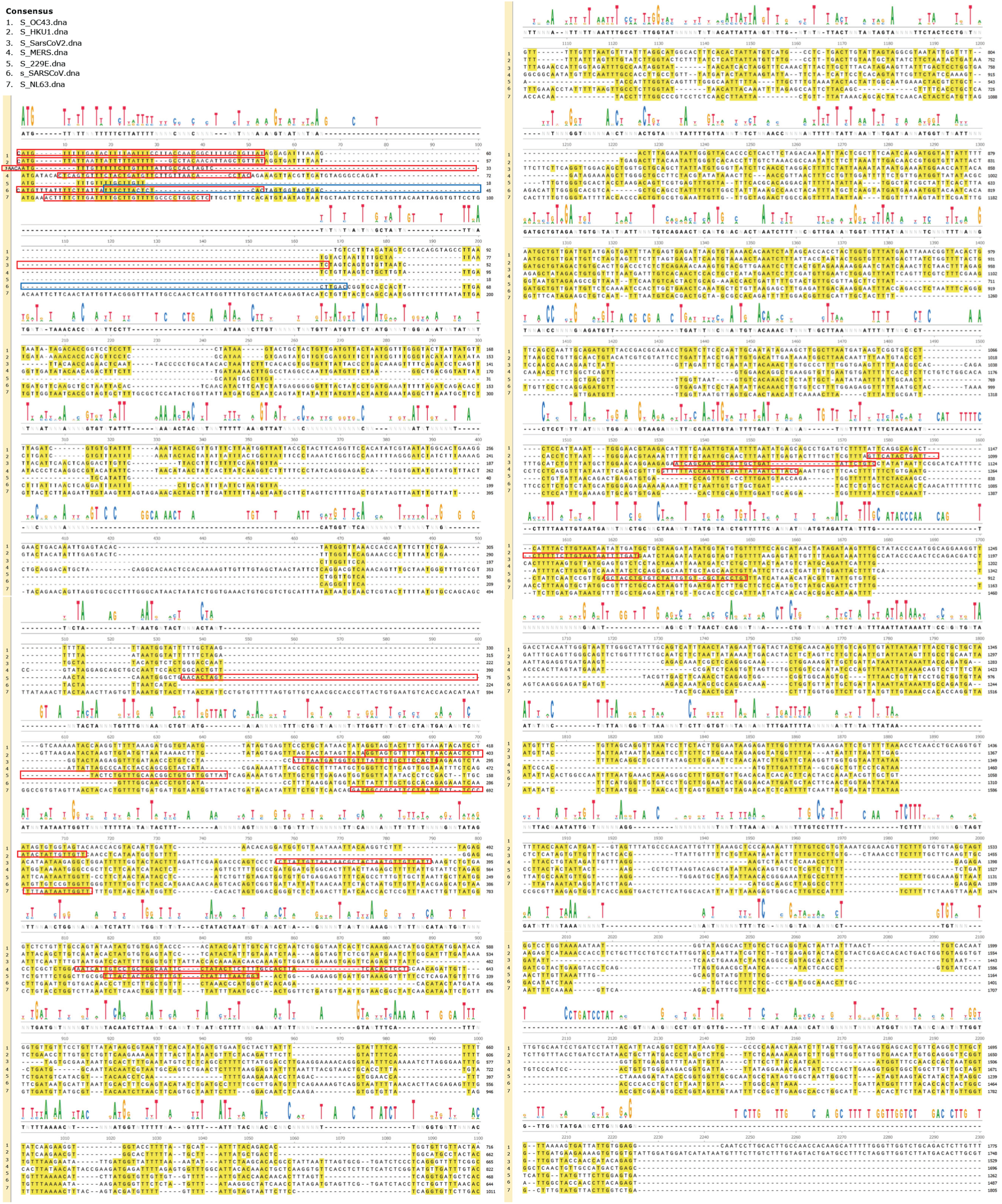
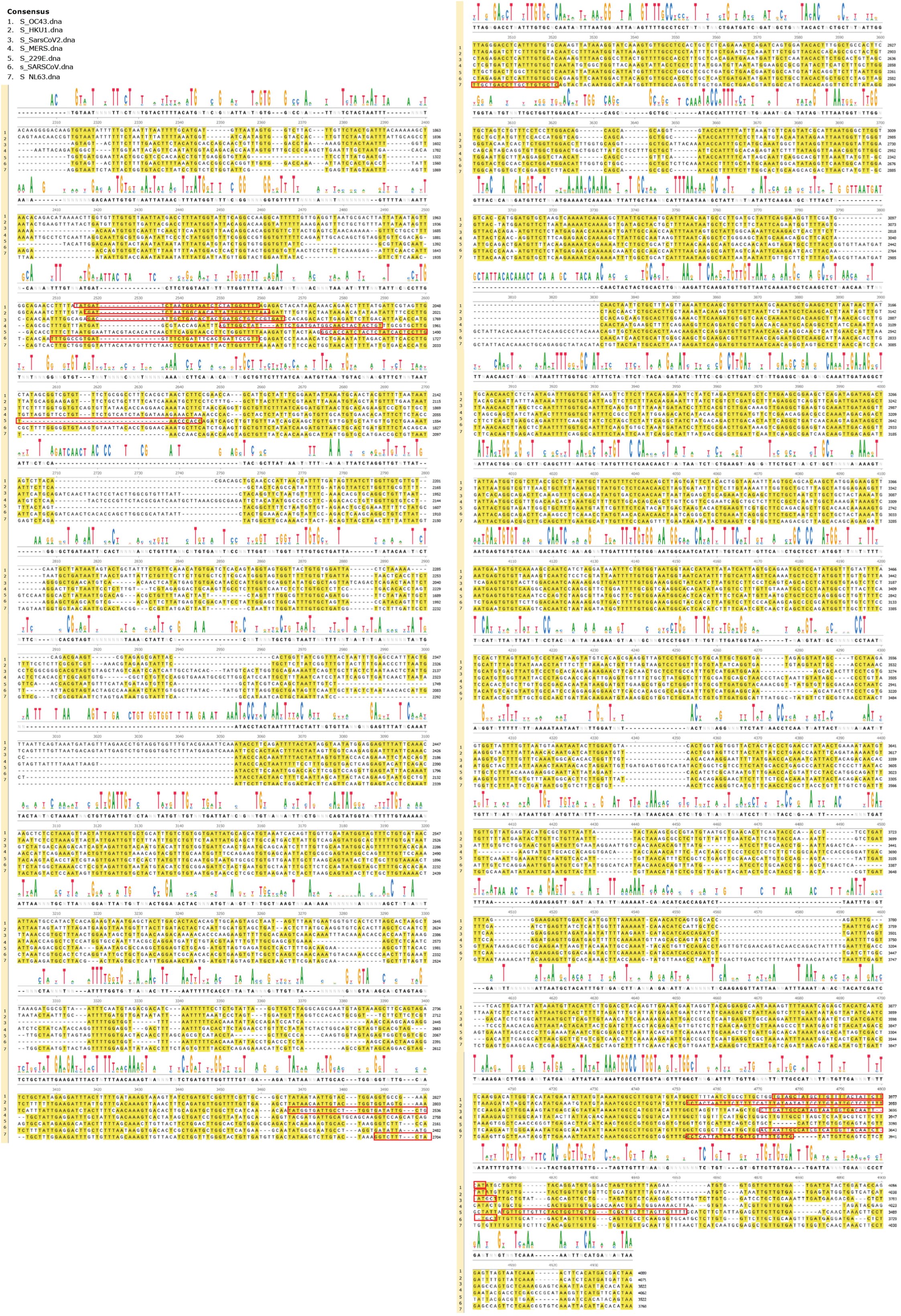
Alignment of S gene from the seven hCoVs. The alignments depict the gene sequence. Red boxes – SECReTE sequences. Blue box – overlapping SECReTE sequence. Yellow highlights – matched bases to the consensus sequence.

**Supplementary Figure S3.**
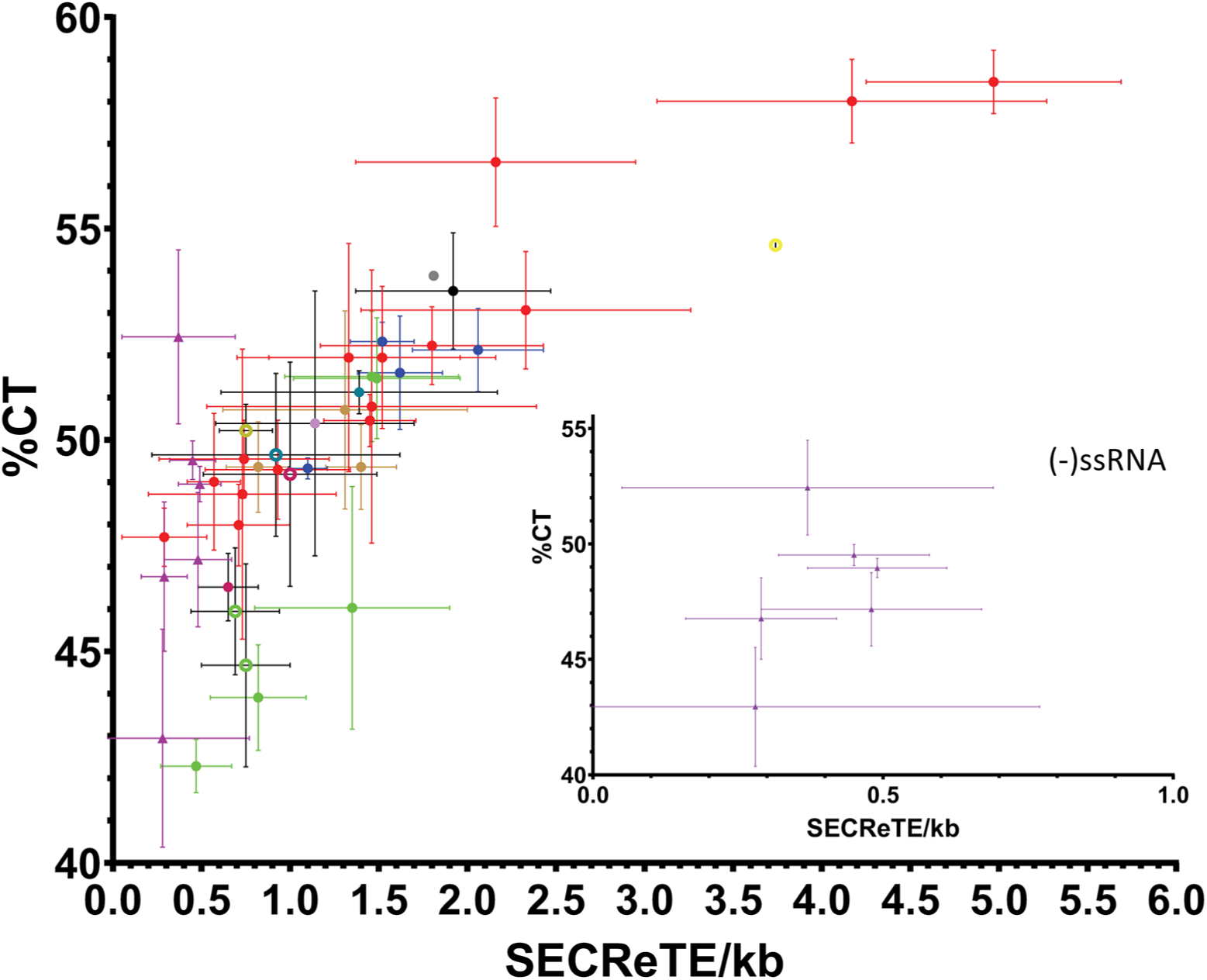
SECReTE elements in (+)ssRNA and (−)ssRNA viruses. The distribution of the average (±S.D.) SECReTE score of each family/genus (color coded) vs the average (±S.D.) percentage of pyrimidines (%CT) in the (+)ssRNA genomes. Inset – same shown for (−)ssRNA viruses only. See also Figure 3 and Supplementary Tables S5 & S6.

**Supplementary Figure S4:**
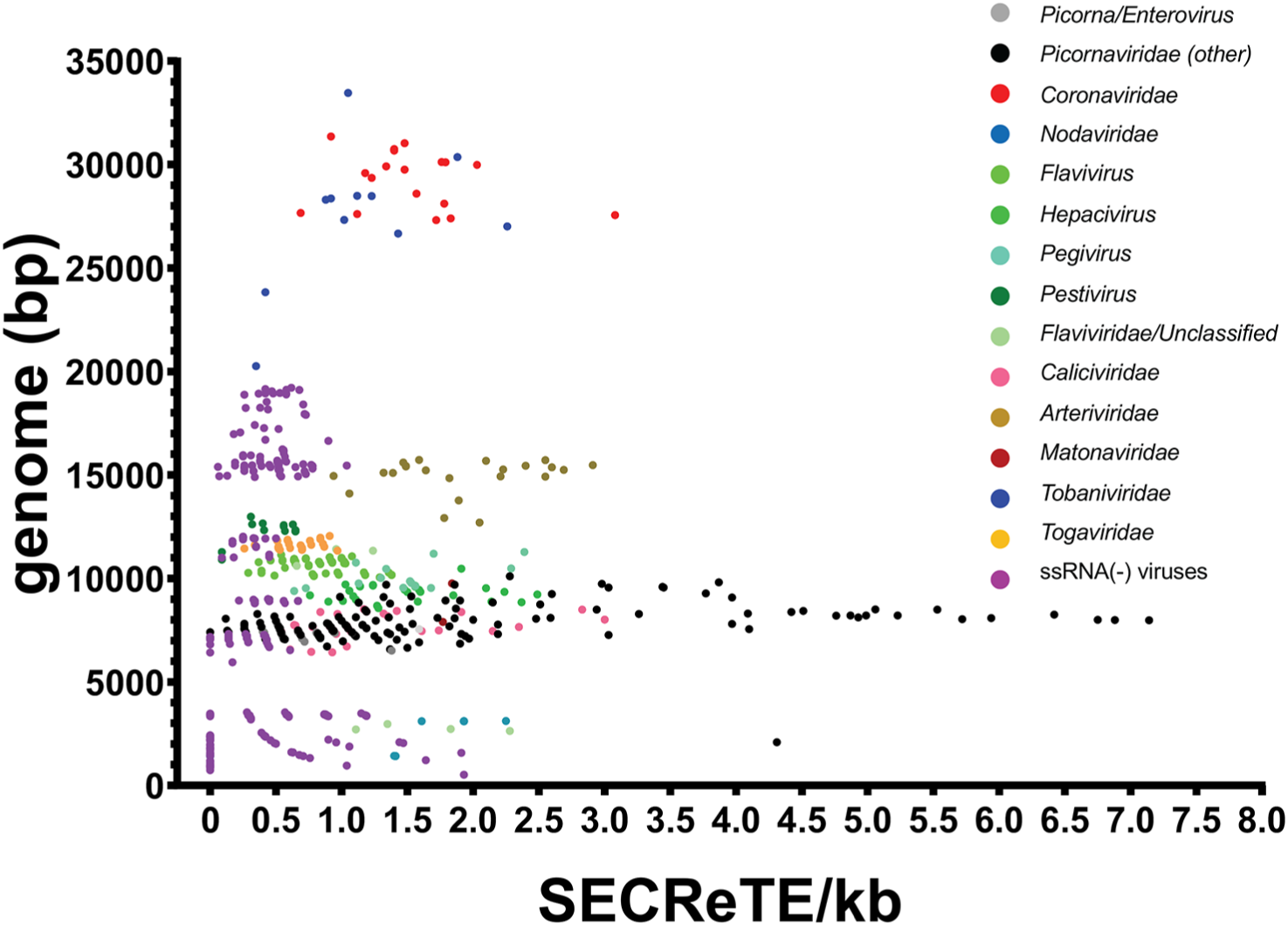
SECReTE elements in (+)ssRNA and (−)ssRNA viruses. The distribution of the SECReTE score of each individual genome (color coded by family/genus) *vs.* the genome size is depicted. See also Figure 3 and Supplementary Tables S5 & S6.

**Supplementary Table 1. List of SECReTEs in the seven human coronaviruses**

Each sheet provides data of the indicated hCoV. *#* – the SECReTE number (as annotated in Figure 1A); *Name –* Accession number in GenBank; *Triplets –* number of triplet repeats in the motif; *Start –* the start location in the viral genome; *Seq –* the SECReTE sequence; *Protein –* the viral protein (or UTR) in which the SECReTE resides; ^ – the frame of the SECReTE relative to the reading frame; *Membrane –* whether the protein a membranal proteins (yes/no); *TMD –* whether SECReTE occurs within a TMD-coding sequence.

**Supplementary Table 2. List of SECReTEs in SARS-CoV-2 strains**

Sheet “sars-cov-2 strains” provides strain accession number in GeneBank (*Name*), genome length (*seqLen*), and number of SECReTEs (SECReTEs). Sheet “SECReTE_seq” provides the SECReTE sequences in the SARS-CoV-2 strains. *Name –* Accession number in GenBank; *Triplets –* number of triplet repeats in the motif; *Start –* the start location in the viral genome; *Seq –* the SECReTE sequence.

**Supplementary Table S3. List of SECReTE in human secretome genes**

SECReTE data for human secretome protein-encoding genes. Sheet “Human secretome” lists the human secretome protein-encoding genes: *Symbol –* Gene symbol; *SECReTE count –* number of SECReTE motifs; *Displayed protein* – protein encoded; *RefSeq –* NCBI reference sequences; *5’UTR –* length of 5’UTR (nucleotides); *CDS –* length of coding region (nucleotides); *3’UTR –* length of 3’UTR (nucleotides); *RefSeqType –* canonical or one isoform; *Signal peptide –* position of signal peptide *TMDs –* position of TMDs; *SECReTE/kb –* SECReTE score per kb. Sheet “SECReTE_seq” provides the SECReTE sequences: *SECReTE ID* – NCBI reference sequence number with start position of motif; *Gene ID* – NCBI reference sequence number, *Symbol –* gene symbol, *Start –* start position of SECReTE; *End –* end position of SECReTE; *# triplets –* number of triplet nucleotide repeats in the motif; *Location* – location in gene; *SigPep/TMD –* SECReTE presence in signal peptide or TMD; *Frame –* reading frame of motif (if applicable), *SECReTESeq –* SECReTE sequence. Sheet “OR-DEF-MUC” provides the data used in Figure 4. *Gene –* gene name, *SEC/kb –* SECReTE number normalized for gene length, *Length –* gene length.

**Supplementary Table S4. List of SECReTEs in Coronaviridae**

Sheet “SECReTE count” summarizes the data for C*oronaviridae*. *Accession –*Accession number in GenBank, *Description –* type of CoV and source, *Closest strain –* according to ^18^, *Genus –* genus of virus, *seqLen –* viral genome length, *SECReTE –* number of SECReTE motifs, *SEC/kb –* SECReTE number per kb, *%CT –* percentage of pyrimidines in genome. Columns K-O summarize data for the CoV genera. Sheet “SECReTE_seq” provides the SECReTE sequences of the strains. See also Figure 2. *Name –* GenBank name, *triplets –* number of consecutive triplet nucleotide repeats in the motif, *Start –* start position of motif in sequence, *Seq –* motif sequence.

**Supplementary Table S5. A list of SECReTEs in ssRNA viruses**

Sheet “Vertebrates” – a list of viruses that infect vertebrates or use invertebrates as vector to infect vertebrates. Sheet “invertebrates” – a list of viruses that infect invertebrates. *Accession –* Accession number in GenBank, *RNA strand –* type of ssRNA, *Family, Genus, Subgenus, Name* – taxonomy for each given virus, *Segment –* segment scored, *Host –* whether host is solely vertebrate or uses invertebrates as vectors, *seqLen –* viral genome length, *SECReTE –* total number of SECReTEs in the genome. *SEC/kb –* overall total SECReTE score, *%CT* – percentage of pyrimidines in genome, *SECReTE(CDS) –* SECReTE number in the coding regions, *SEC(CDS)/kb –* SECReTE score per kb in coding region (Data obtained from Supplementary Table S6). Sheet “averages” provide average data and biological information on viral replication centers for specific families/genera. *Avg. SECReTE –* average number of SECReTEs(CDS) per kb, *SD –* standard deviation, *Avg %CT –* average pyrimidine content, *Order, Family/Genera –* taxonomy of virus, *Viruses –* specific viruses in which VRCs were studied, *Organelles of Replication –* membrane source, if known, for viral replication, *Membrane structure –* types of membranes generated for viral replication. *SEC(CDS)/kb* – the SECReTE scoring used in Figure 3, Supplementary Figures S3 and S4, and Table 2.

**Supplementary Table S6. List of all SECReTEs in ssRNA viruses**

Sheet “all SECReTE” provides all SECReTE sequences in the ssRNA genomes. *Genome –* NCBI RefSeq genome identity number, *Start –* start position of SECReTE in genome, *Length –* number of nucleotides, *Sequence –* SECReTE sequence. Sheet “SECReTE in genes” provides the sequences of SECReTE in coding sequences. *Genome* – viral genome, *Gene –* number of gene in genome, relative to gene order, *Location –* location of SECReTE in genome, *Length –* number of nucleotides, *Sequence –* SECReTE sequence. Sheet “non-redundant” – same as “SECReTE in genes” however all redundant (*i.e.* overlapping) SECReTEs removed. Sheet “SECReTE count” – lists the numbers of SECReTEs *(#SECReTE*) used in “SECReTE(CDS)” shown in Supplementary Table S5.

## Notes

### Competing Interest Statement

The authors have declared no competing interest.

